# The impact of intermittent palatable food consumption on microbiota structure in male and female rats

**DOI:** 10.64898/2026.02.12.705550

**Authors:** Sarah Gore, Ahlam Akmel, Saadiya Jackson, Alexa Ryan, Haley Warren, Courtney Robinson, Kimberlei Richardson

## Abstract

The gut microbiome plays a vital role in metabolism, behavior, and overall health, with diet being a key factor shaping its composition. This study examines the impact of intermittent palatable food (PF) consumption on microbiota structure in male and female rats, focusing on feeding preferences and sex differences. Rats were characterized as high preferring (HP) or low preferring (LP) based on PF intake, and microbial analyses were conducted across different gastrointestinal regions, including the colon, feces, cecum, and cecal contents. However, microbiota composition varied with significant differences in Firmicutes, Bacteroidetes, and Actinobacteria abundances. Sex-based differences were evident particularly in fecal and cecal samples, where Proteobacteria and Actinobacteria populations varied between males and females within the same feeding groups. Our findings support the notion that dietary habits and microbiota composition may form a feedback loop, reinforcing food preferences through gut-brain axis signaling. While alpha diversity remained unchanged, beta diversity analysis indicated subtle, but significant differences in microbial community structures based on sex and feeding behavior. This research provides novel insights into the interplay between diet, gut microbiota, and behavior, while emphasizing the importance of considering sex as a variable in microbiome studies. Understanding these relationships may inform dietary interventions aimed at optimizing microbiota composition to improve metabolic and mental health outcomes linked to diet-induced microbiota shifts.

**Importance:** The gut microbiome plays a critical role in various bodily functions, from the brain’s protective mechanisms to dietary behaviors and food choices. In this study, we sought to broaden the understanding of how various levels of high fat and high carbohydrate diet consumption alter gut microbiota, and ultimately, shape food preference behaviors. We assessed preference behaviors by categorizing rats into high-preference and low-preference groups based on their consumption of high calorie, palatable food, then analyzed their gut bacterial composition, comparing diet preference groups and examining whether sex differences were reflected in the results. The findings from this research hold significant implications for understanding the complex interplay between diet, microbiota, and behavior. Understanding the bidirectional relationship between feeding behavior and microbiota may provide novel insights into the mechanisms underlying eating disorders and metabolic dysregulation.

## Introduction

The gut microbiome is an ecosystem of trillions of microorganisms that plays a key role in host physiology, influencing various bodily functions. The relationship between diet and the microbiota is particularly significant, as dietary choices can profoundly alter microbial composition and host health. For example, the blood-brain barrier permeability is influenced by these microbes (1) as is neurotransmission (2) and host behavior (3,4) including diet choice and consumption (5,6). Host diet is among the most influential environmental factors connected to the composition and structure of this microbial community (7-10). Diversity in a host’s diet can result in many metabolic outcomes with an assortment of benefits. Macronutrients such as carbohydrates, fats, and proteins are catabolized by gut microbes to short chain fatty acids (SCFA), including acetate, propionate, and butyrate. They impact host physiology by being further metabolized by host cells, for example as energy substrates, and/or functioning as regulatory signaling molecules (11). Interestingly, a host seeking particular diets and the microbiota associated with the host may be part of a positive feedback loop. This would allow certain bacteria to thrive on specific energy sources from the host diet and generate products that can be used by the host (e.g. SCFA). This phenomenon leads to the host preferentially feeding on those diets thereby selecting the gut microbiota that yield products of interest for the host (10). An emerging area of interest is the impact of intermittent palatable food consumption on gut microbiota structure, especially in the context of macronutrient intake and host behavior.

Macronutrients such as carbohydrates and fats are vital to many animal diets (12) and dietary modulation of gut microbiota. The influence of various carbohydrate and fat compositions on the gut microbiota can range from improving prebiotic health and metabolism to causing disruptive metabolic dysfunction (13). The intake of fiber, a complex carbohydrate, has a positive effect on gut microorganisms by increasing the population of microbes that metabolize certain carbohydrates to create SCFA and have positive impacts on overall host health (14). On the contrary, high levels of certain other carbohydrates, such as fructose, can have detrimental effects on the gut microbial environment and increase intestinal mucosal inflammation, disturbance on the epithelial barrier, and rearrangement of the catabolic products from the microbiota (15). Fats are very similar in that various fatty acids cause different changes to the gut microbiome and behavior. In general, high consumption of dietary fats results in gut dysbiosis, or the loss of beneficial members of the microbiome, which together are associated with systemic inflammation, diabetes, and cardiovascular disease, among other metabolic disorders and chronic diseases (16-19). In addition, it has been proposed that high-fat diets can lead to changes in behavior. For example, rats fed a high-fat diet displayed increased motivation to consume more sucrose compared to chow-fed only rats (20). More recent work has implicated changes to the microbiota-gut-brain axis as a result of high-fat diets being a driver of anxiety-like behavior in rats (21).

Another factor that can influence the gut microbiome is sex and it may differentially impact the microbiota (22,23). It is possible that sex-diet interaction can regulate gut microbiota and generalized assumptions about the effects of food choice on microbiota composition should be avoided (24). More studies are needed to determine how sex differences play a role in food choice and how sex-diet interactions affect microbial communities.

Intermittent feeding models provide a unique perspective on how dietary choices shape microbiota over time. Many studies have focused on various feeding models to better delineate the underlying mechanisms that control the microbiome and its impact on feeding behavior. The intermittent feeding model can be used to explore the impact of chronic high fat and sugar on food preference behavior (25). Intermittent consumption of high fat and high carbohydrate diets has been linked to metabolic and neurobiological changes. Yet the specific interactions between such feeding patterns and gut microbiota remain underexplored. We seek to expand the use of this model and identify how sex and changes in preference can shift microbial communities. In this study, high-preferring (HP) and low-preferring (LP) phenotypes emerged using the intermittent PF exposure, reflecting inherent individual differences in food preference. HP rats consistently consume more PF when available, whereas LP rats exhibit greater reliance on standard chow. We sought to provide a greater understanding of how varied consumption of high fat and high carbohydrate diets can impact the gut microbiota and how that may influence food preference behaviors (26). We compared the bacterial communities in the colon, cecum, cecum contents and fecal samples between males and females, as well as each of the feeding preference groups. Using this approach, we investigated the influence of innate preference on how and what an animal seeks as food, how changes in diet composition can shape the gut microbiota, whether sex differences are reflected in these outcomes. Intermittent PF consumption represents a compelling model for studying the dynamic relationship between diet and gut microbiota. By integrating behavioral phenotyping, microbial analysis, and sex-based comparisons, this research contributes to a deeper understanding of how dietary habits shape gut microbial ecology and influence host health. Through these insights, future studies may explore targeted dietary interventions aimed at optimizing microbiota composition and improving metabolic outcomes.

## Methods

### Animal/Study Population

Adult female and male Sprague-Dawley rats (200-250 g, n= 57 rats, Charles River, Raleigh, NC, USA) were housed individually and maintained on a 12-hour reverse light-dark cycle. They were provided standard chow (7012 LM-485, Teklad Diets, Madison, WI, USA) and water *ad libitum* throughout the entire study.

### Feeding Test to Determine Eating Phenotype

The weights of the rats and standard chow were recorded daily. On feeding test days, rats (n=48) had *ad libitum* access to standard chow and 30 grams of high fat, sweet palatable food (PF) pellets (D12451, 45% energy from fat, 20% kcal from protein, 35% kcal from sugar; Research Diets, Inc., New Brunswick, NJ, USA) for four hours. The weights of the standard chow, PF, and each rat were recorded 1 and 4 hours after the PF was introduced. The PF was removed after 4 hours. The 4-hour time period was used to resemble the interval of consumption that is associated with binge eating in humans (27). The weights of the rats and standard chow were also recorded the following day (24 hours after PF introduction). Rats were provided only standard chow for 2-3 days between each feeding test.

The amount of PF recorded at the 4-hour time point was used to assign the rats to their respective preference categories (28). Rats with median PF intake in the upper tertile for at least 50% of the feeding tests were classified as high preferring (HP; n=12), and those with PF intake in the lower tertile for at least 50% of the feeding tests were classified as low preferring (LP; n=10). The remaining rats were classified as neutral and were not used further in this study. A total of eight intermittent feeding tests were conducted to determine whether the rats demonstrated a high preferring, low preferring, or neutral feeding phenotype associated with PF intake. Rats (n=9) that did not participate in the PF feeding tests and only received *ad libitum* access to standard chow and water were classified as chow-only control rats. Following the final feeding test, the rats were deeply anesthetized with ketamine/xylazine (80/10 mg/kg, i.p.) and cecum, cecum contents, colon and fecal pellets were extracted and quick frozen until processing for DNA extraction.

### DNA extraction

Total DNA was extracted using the Invitrogen PureLink® Microbiome DNA Purification Kit (ThermoFisher) following the manufacturer’s instructions, using a Mini-BeadBeater 16 (BioSpec Products) for homogenization of samples. Eluted DNA was stored at -20^°^C.

### 16S rRNA gene sequencing

The V4 region of the 16S rRNA gene was amplified from each sample using the dual-indexing sequencing strategy developed by Kozich et al (29). Amplification and sequencing was conducted by the Center for Microbial Systems at the University of Michigan on the Illumina MiSeq platform, using a MiSeq Reagent Kit V2 500 cycles or a MiSeq Reagent Nano Kit V2 300 cycles, according to the manufacturer’s instructions with modifications as described previously (29-31).When necessary due to low biomass, a “touchdown PCR” protocol was followed: one cycle of 95^°^C for 2min followed by 20 cycles of 95^°^C for 20s, 60^°^C for 15s, and 72^°^C for 5 min with a decrease of 0.3^°^C each cycle, followed by 20 cycles of 95^°^C for 20s, 55^°^C for 15s and 72^°^C for 5 min, and a final extension of 72^°^C for 10m Concentrations of libraries and pool normalizations were determined conducted using the Life Technologies SequalPrep Normalization Plate and the Kapa Biosystems Library Quantification kits as previously reported (29-31). Amplicons of the appropriate size were extracted from agarose gels using the QIAQuick Gel Extraction kit (Qiagen). The final library consisted of equal molar amounts of amplicons from each of four plates, normalized to the lowest concentration of the four. When a library concentration was <1.0nM, the protocol of Quail et al was used for denaturation (32). The final load concentrations were 4 – 5.5 pM with a 10-15% PhiX spike added to increase diversity. Sequencing reagents were prepared as described further on the SchlossLab website (33) and custom read 1, read 2, and index primers were added to the reagent cartridge (29-31). Finally, FASTQ files were generated for paired-end reads.

### Community Analysis

Sequences were processed and analyzed using mothur v1.48.0 using the MiSeq SOP described at (33). After paired-end sequence reads were assembled and short sequences and/or sequences containing ambiguous bases were removed, sequences were aligned and screened for chimeras. The remaining sequences were then classified using a mothur-formatted Ribosomal Database Program training set (v.9; minimum confidence score of 80%) (34). All sequences that are classified as chloroplast, mitochondria, eukaryote, or unknown at the Kingdom taxonomic level were removed. Operational taxonomic units (OTUs) were generated by binning sequences that were at least 97% similar using the Opticlust method (30,35). The taxonomic affiliation was then determined for each OTU using the supplied classification or when the supplied classification was inconclusive, sequences were compared to NCBI nr database using BLAST (36). Community richness was assessed using both observed richness and the Chao-1 richness estimator calculated using OTU-based analyses (37). Community diversity was measured using the Inverse Simpson index. Principal component analyses were conducted on the Bray-Curtis dissimilarity distances (38). Data were visualized and statistically significant differences were determined using ANOVA followed by post-tests as needed using standard R packages (39-41). Differences in community structure between treatment groups and sample types were assessed with analysis of molecular variance (AMOVA) and Metastats in mothur (33,42).

## Results

### Diet Preference Feeding Tests

In order to determine the food preferences of the rats, multiple tests were conducted to measure the amount of PF and standard chow consumed on each feeding test day. Female and male HP rats consumed significantly more PF than female and male LP rats, respectively, one hour after the beginning of the feeding test (Figure 1A, *p<0.0001, ^#^p<0.05). When PF intake was evaluated with respect to the average weights of each group, the female HP rats still consumed significantly more than female LP rats (Figure 1B, ^*^p<0.05). There was no significant difference in PF intake in male rats at the 1hour timepoint when consumption was standardized to the average weights of each group (Figure 1B, p=0.0960). Similarly, female and male HP rats consumed significantly more PF than female and male LP rats, respectively, four hours after the beginning of the feeding test (Figure 1C, *p<0.001, ^#^p<0.01). When the PF intake was evaluated with respect to the average weights of each group, the female HP rats still consumed significantly more than female LP rats (Figure 1D, ^*^p<0.05) at the four-hour interval. The same significant pattern was noted when comparing male HP and male LP rats (Figure 1D, *p<0.05).

**Figure 1.**
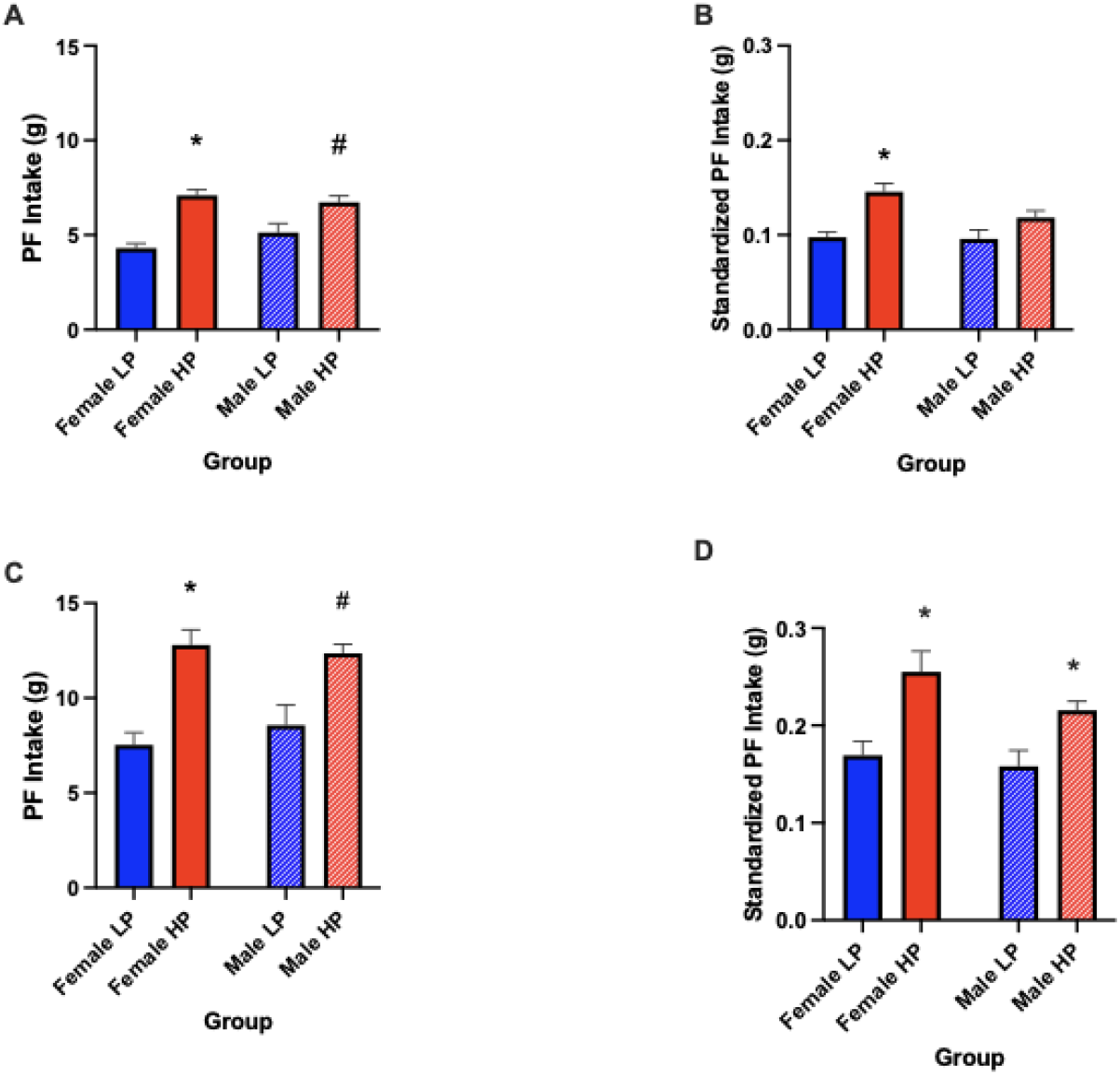
These graphs represent the average amount of PF consumed by male and female LP and HP rats at different time points during the intermittent feeding tests. 1A) One hour after the start of the tests, female and male HP rats ate significantly more PF than LP rats female and male LP rats (*p <0.0001), ^#^p <0.05), 1B) After PF intake was standardized based on rat weights, female HP rats still consumed significantly more PF than female LP rats one hour after the start of the tests (*p <0.05), 1C) Four hours after the start of the tests, HP rats continued to consume significantly more PF than LP rats for female and male subjects (*p <0.001, ^#^p <0.01) and 1D) After four hour PF intake was standardized based on rat weights, female and male HP rats still consumed significantly more PF than female and male LP rats, respectively (*p <0.05).

Normal chow intake was also recorded for all rat groups during the feeding tests. As a baseline for regular standard chow intake, the chow only group consumed standard chow throughout the entire study. It was noted that the female chow only group consumed more chow on the feeding test days than both the female LP and HP groups (Figure 2A, F (2, 16) = 95.68 *p<0.0001, #p respectively). Also, female LP rats consumed significantly more standard chow than HP rats 24hours after the start of the feeding tests (Figure 2A, ^^^p<0.001). Likewise, the male chow only group consumed more chow on the feeding test days than male HP rats, but there was no significant difference when compared with standard chow intake for the LP group (Figure 2A, *p<0.05, p=0.3663, respectively). These data were also standardized with respect to the weights of the rats (Figure 2B). When male and female data were combined for chow-only, LP and HP groups and the means were compared for standardized chow intake 24 hours after the feeding tests, there was a significant difference between the groups (Figure 2B, F (2, 28) = 70.51, *p<0.0001). In Figure 2C, the chow-only (male and female data combined) group consumed significantly more standard chow than both groups with prior PF exposure (*p<0.0001) and the combined-sex, LP group consumed more standard chow than the combined-sex, HP group (**p<0.001).

**Figure 2.**
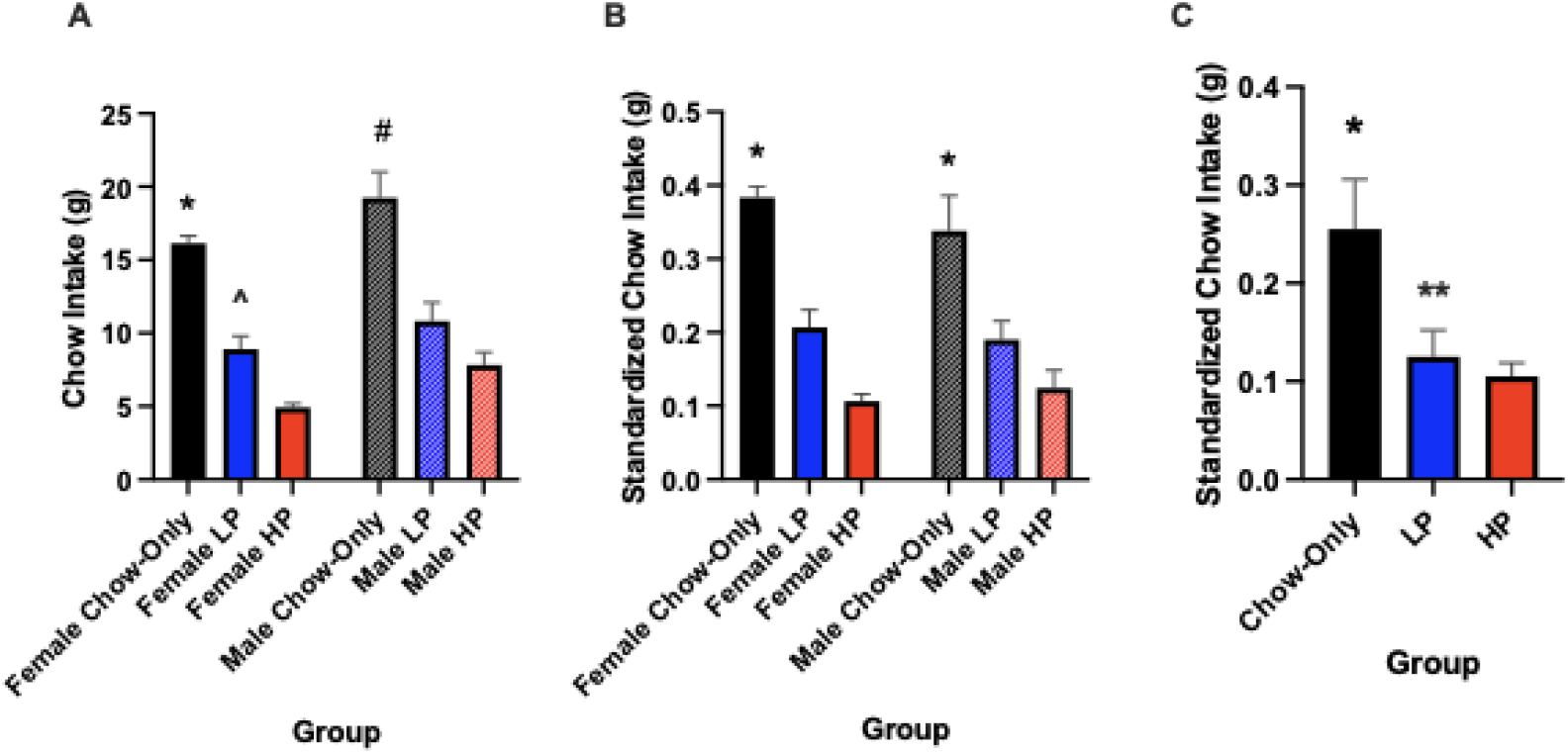
The data represented above are for total standard chow intake 24hours after the start of the feeding tests. 2A) Female and male chow only rats consumed significantly more standard chow on feeding test days versus LP and HP rats. The female LP rats consumed more standard chow than female HP rats. (*p<0.0001, ^^^p<0.001, ^#^p<0.05). 2B) This graph represents the standardized data with respect to weight for the 24-hour intake of standard chow consumed by HP and LP rats on feeding test days, in comparison with the chow only groups (female chow vs LP and HP*p<0.0001); male chow vs. LP and HP, *p<0.0001). 2C) Data from female and male groups were combined for each group, and the chow-only group had a significant increase in standardized chow intake versus the LP and HP groups. (*p<0.0001, ^p <0.001). (The data represented by the mean + SE, n=31.)

Data from the consumption of standard chow were compared for HP, LP and Chow-only control rats on non-feeding test days. There were no significant differences in standard chow consumption when comparing HP, LP and chow-only, control rats (Figure 3A, females: F (2, 16) = 2.234, p=0.1395, males: F (2, 9) = 1.680, p= 0.2399). The data were standardized with respect to weight, and no statistical differences were observed (Figure 3B). So, on days when all groups received the same diet (the standard chow), there were no observable differences in rats that received PF versus non-PF diet rats. Significant differences were observed when male and female data were combined revealing that the combined chow only group consumed more standard chow than both groups with prior exposure to PF (Figure 3C, F (2, 27) = 4.441, *p<0.05).

**Figure 3.**
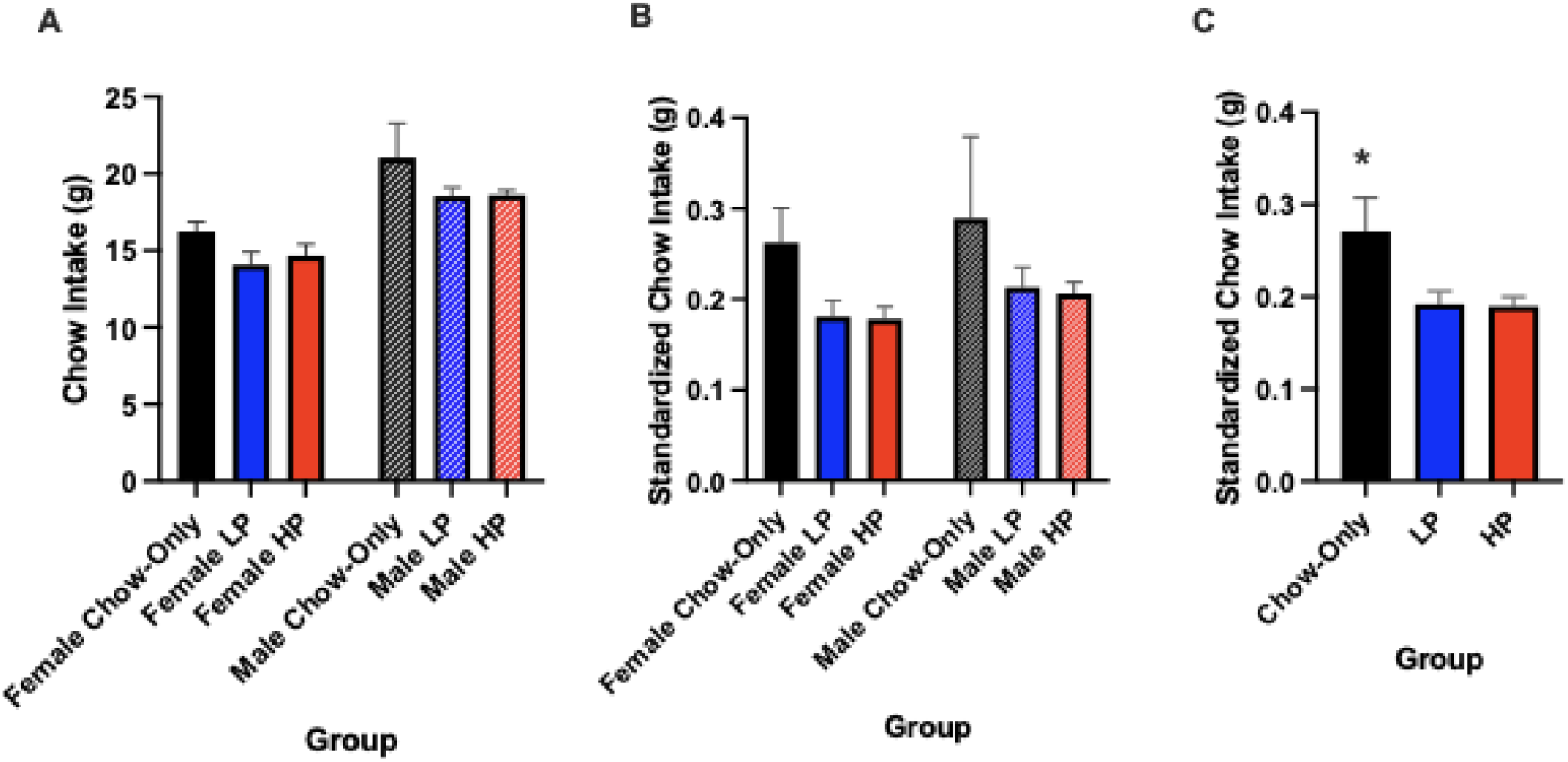
The graphs above show the amount of standard chow consumed on non-feeding test days by all three rat groups: chow only, HP, and LP. 3A) The data represent the unstandardized chow intake data and 3B) The standardized data are presented with respect to the weight of the rats. 3C) Significant differences were observed when male and female data were combined revealing that the combined chow only group consumed more standard chow than both groups with prior exposure to PF (*p<0.02). Data are represented as mean + SE, n=31.

Observations were also noted for changes in bodyweight during the feeding tests. While there were differences in the amount of PF consumed, no significant differences in body weights were noted between the HP and LP groups at the 4-hour timepoint for females (Figure 4A, F (2, 16) = 1.424 p = 0.2696) and males (Figure A, F (2, 9) = 0.5393, p=0.6009). Similarly, there were no significant differences in body weights at the 24-hour timepoint for females (Figure 4B, F (2, 16) = 0.9577, p = 0.4047) and males (F (2, 9) = 0.3550, p=0.7106). The data in Figure 4C represent male and female data combined for chow only, LP and HP groups, and no significant differences were noted. (Figure 4C, F (2, 28) = 0.4404, p=0.6482). Likewise, there were no significant differences in 24-hour bodyweight on non-feeding test days when comparing combined sex-dependent data for chow only, LP and HP rats F (2, 28) = 0.7629, p=0.4757.

**Figure 4.**
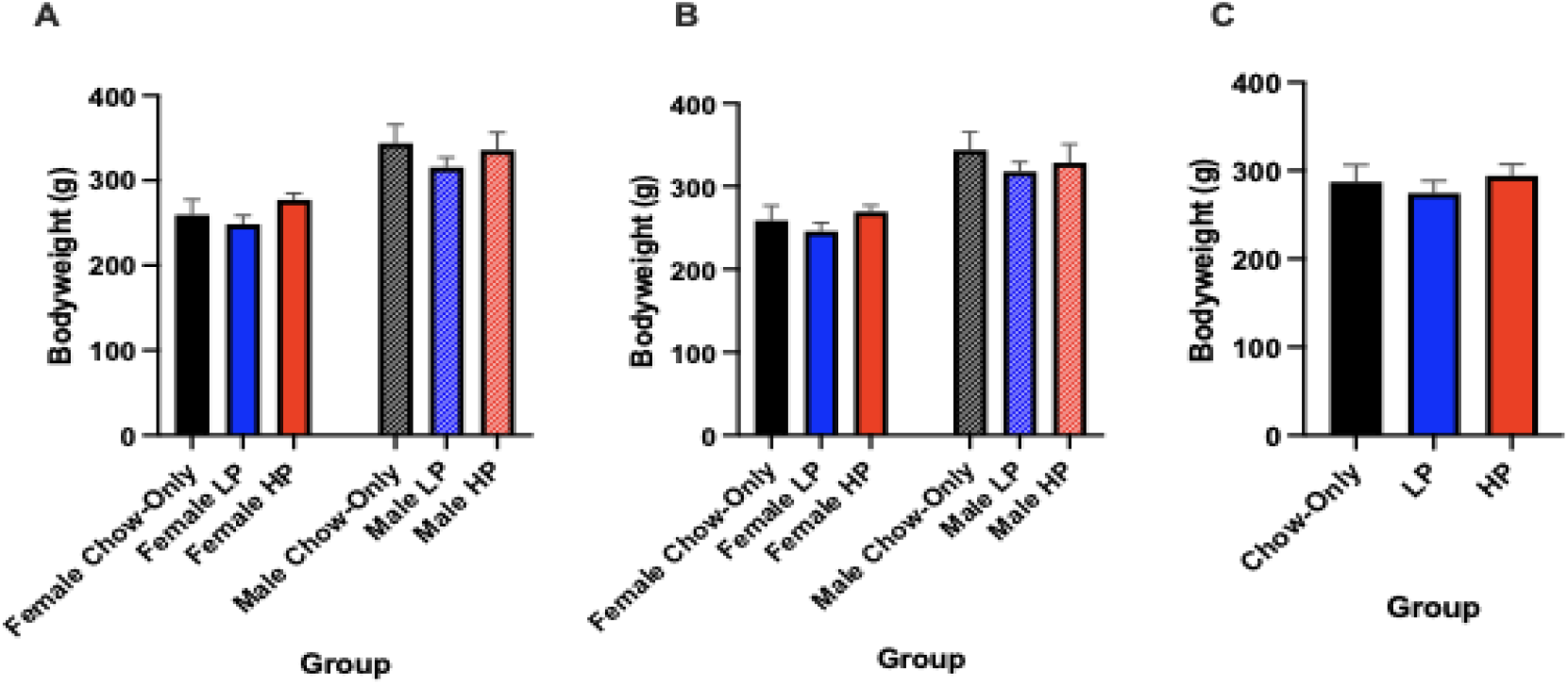
Body weights of the HP and LP rats at different time points.4A) Weights of rats 4 hours and 4B) 24 hours after the start of the feeding tests. The chow only data are included for comparison with the groups with prior PF exposure. There were no significant differences in bodyweights between the groups at either timepoint. 4C) Even when the 24-hour data for male and female groups were combined, there was no significant difference in body weights. Data are represented as mean + SE, n=31.

### Characteristics of the Bacterial Community-Alpha Diversity, Composition, and Structure

#### Alpha Diversity Analysis

Microbial community richness and diversity measures were calculated using OTU-based analyses. Inverse Simpson index reflects community diversity while Observed Richness and Chao index reflect the species’ richness. When comparing the control baseline samples, there were no significant differences observed in all of the alpha diversity measures as shown in **Table 1**. To determine significant differences in the diversity measures, ANOVA test was performed on samples with normal distribution, and Kruskal-Wallis test was performed for samples that did not have a normal distribution (Shapiro Wilk test was used to test for normal distribution). According to the ANOVA and Kruskal-Wallis test, there were no significant differences (p<0.05) among the control baseline feeding preference groups. Therefore, with respect to richness and evenness, the microbial structure within the different feeding groups did not show change in species diversity within the community.

**Table 1:** Alpha Diversity of species richness and evenness Indicated values are the mean values for samples sequenced from each feeding preference group. ANOVA test was performed at a significance level of p<0.05. No significance was observed.

| Feeding Preferences |  | Inverse Simpsons Index | Observed OTUs | Chao1 |
| --- | --- | --- | --- | --- |
| Fecal Sample | **Chow**<br>Female (n=6)<br>Male (n=3) | 21.11<br>22.29 | 198.14<br>208.66 | 331.58<br>324.01 |
|  | **Low-Preferring**<br>Female (n=6)<br>Male (n=4) | 17.32<br>16.92 | 177.714<br>193.36 | 283.31<br>306.24 |
|  | **High-Preferring**<br>Female (n=7)<br>Male (n=5) | 17.32<br>24.19 | 196.74<br>225.41 | 320.14<br>371.34 |
| Colon Sample | **Chow**<br>Female (n=5)<br>Male (n=2) | 17.74<br>29.41 | 96.34<br>119.70 | 149.75<br>199.93 |
|  | **Low-Preferring**<br>Female (n=4)<br>Male (n=3) | 16.04<br>22.04 | 110.68<br>124.74 | 197.17<br>217.12 |
|  | **High-Preferring**<br>Female (n=6)<br>Male (n=4) | 20.90<br>29.37 | 114.78<br>125.32 | 177.83<br>209.25 |
| Cecum Content Sample | **Chow**<br>Female (n=6)<br>Male (n=3) | 33.64<br>32.71 | 185.63<br>172.63 | 313.71<br>295.59 |
|  | **Low-Preferring**<br>Female (n=6)<br>Male (n=4) | 29.02<br>29.95 | 178.01<br>183.97 | 315.94<br>313.80 |
|  | **High-Preferring**<br>Female (n=7)<br>Male (n=5) | 28.89<br>34.86 | 178.15<br>184.66 | 310.78<br>204.30 |
| Cecum Sample | **Chow**<br>Female (n=6)<br>Male (n=3) | 33.20<br>37.04 | 141.53<br>141.51 | 241.03<br>241.12 |
|  | **Low-Preferring**<br>Female (n=5)<br>Male (n=4) | 30.93<br>24.72 | 148.45<br>137.06 | 255.77<br>234.60 |
|  | **High-Preferring**<br>Female (n=7)<br>Male (n=5) | 30.69<br>37.47 | 134.30<br>150.88 | 228.41<br>249.30 |

#### Community Composition and Structure

A principal component analysis was used to understand similarities and variation in the microbial communities and compare the diversity between communities. To understand the differences between communities, beta diversity measures were conducted based on the Bray-Curtis distance matrix.

No significant differences in diversity were detected in alpha diversity across sample types (fecal, colon tissue, cecum content and cecal tissue), rat sex, or feeding behavior group (Table 1 and Figure 5). The predominant phylum identified across all samples was *Firmicutes*, accounting for more than 70% of the sequences analyzed, followed by *Bacteroidetes* (Figure 5 & Figure 6). Differences were detected between samples from male and female rats within the same diet preference group. For example, a significant difference was observed in the *Actinobacteria* abundances in fecal samples from males and females in the chow group (^#^p-value=0.0285) (Figure 5A). In colon samples, there were differences observed at the phylum level between the *Proteobacteria* in male and female chow groups (Figure 5B, ^#^p-value =0.047). Furthermore, a significant difference was detected between the abundances of *Proteobacteria* in the cecum contents of male and female low-preferring groups (p-value = 0.0213). Males in the chow group contained a larger abundance of an unclassified phylum in the cecum contents than females in the same group (p-value = 0.023111).

**Figure 5.**
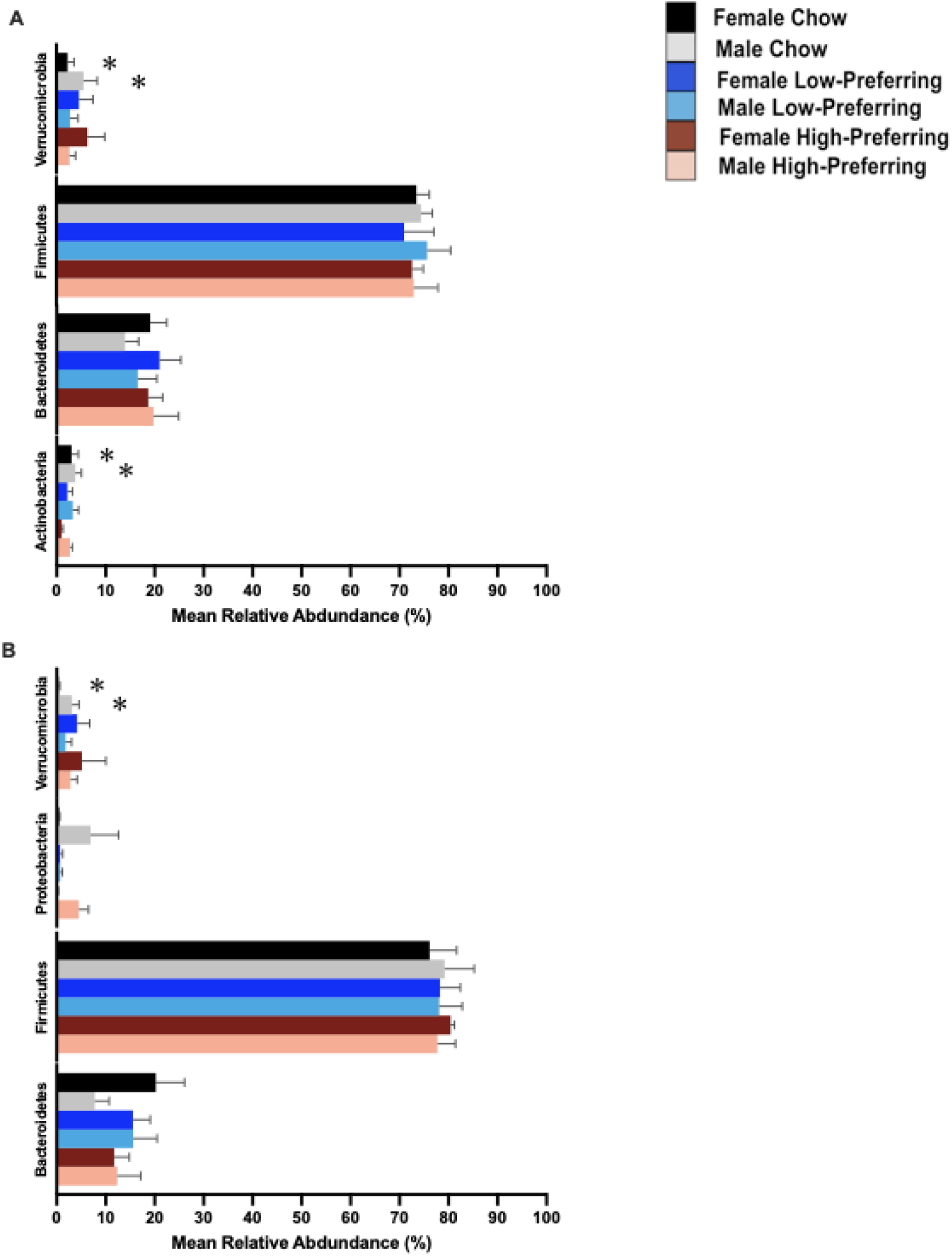
The bacterial phyla distribution represents taxa with >2% relative abundance. Significant differences of p<0.05 between treatment groups are shown using (*). The bar plot distributions are bacterial composition from A) Fecal samples (n=31) and B) Colon samples (n=24).

**Figure 6.**
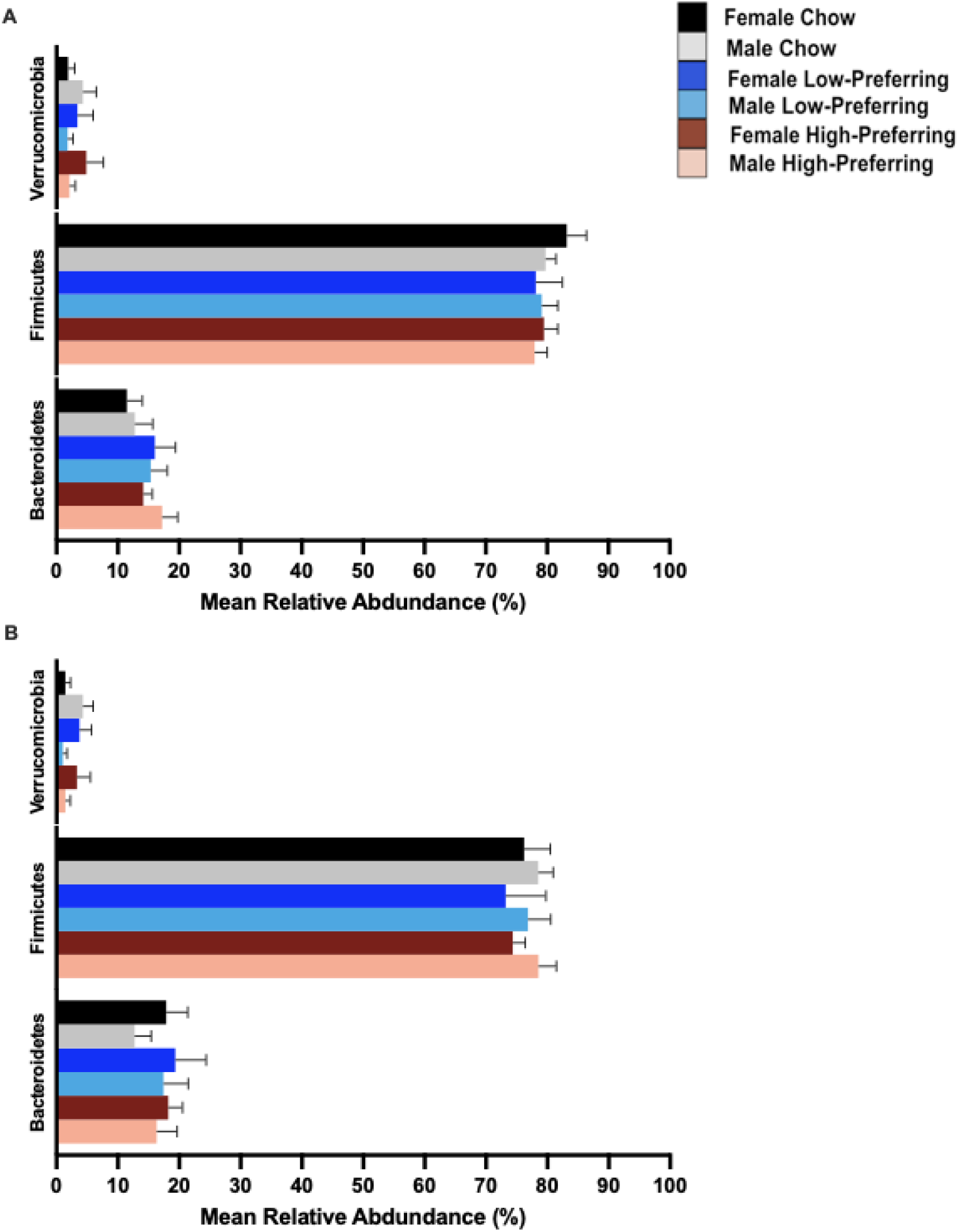
The bacterial phyla distribution represents taxa with >2% relative abundance. The bar plot distributions are bacterial composition from A) Cecum Contents samples (n=31) and B) Cecal samples (n=30). No significant differences were observed between treatment groups.

Significant differences were also observed in feeding behaviors within the same sex. For example, the abundance of the *Actinobacteria* in the colons of female high-preferring rats was higher than in female rats in the chow only group (p-value = 0.029111, see appendix C). Similarly, the cecum content samples in male High-Preferring rats contained more *Actinobacteria* than rats in the chow groups (p-value = 0.034, see appendix F). There were no significant differences detected when structure and beta-diversity were investigated. However, it was noted that members from the *Firmicutes* phyla were responsible for shaping the structure of all communities, *Bacteroidetes* members were influential in the communities in feces and cecal contents, and the *Actinobacteria* were important in the communities in feces and colon (Figure 7).

**Figure 7.**
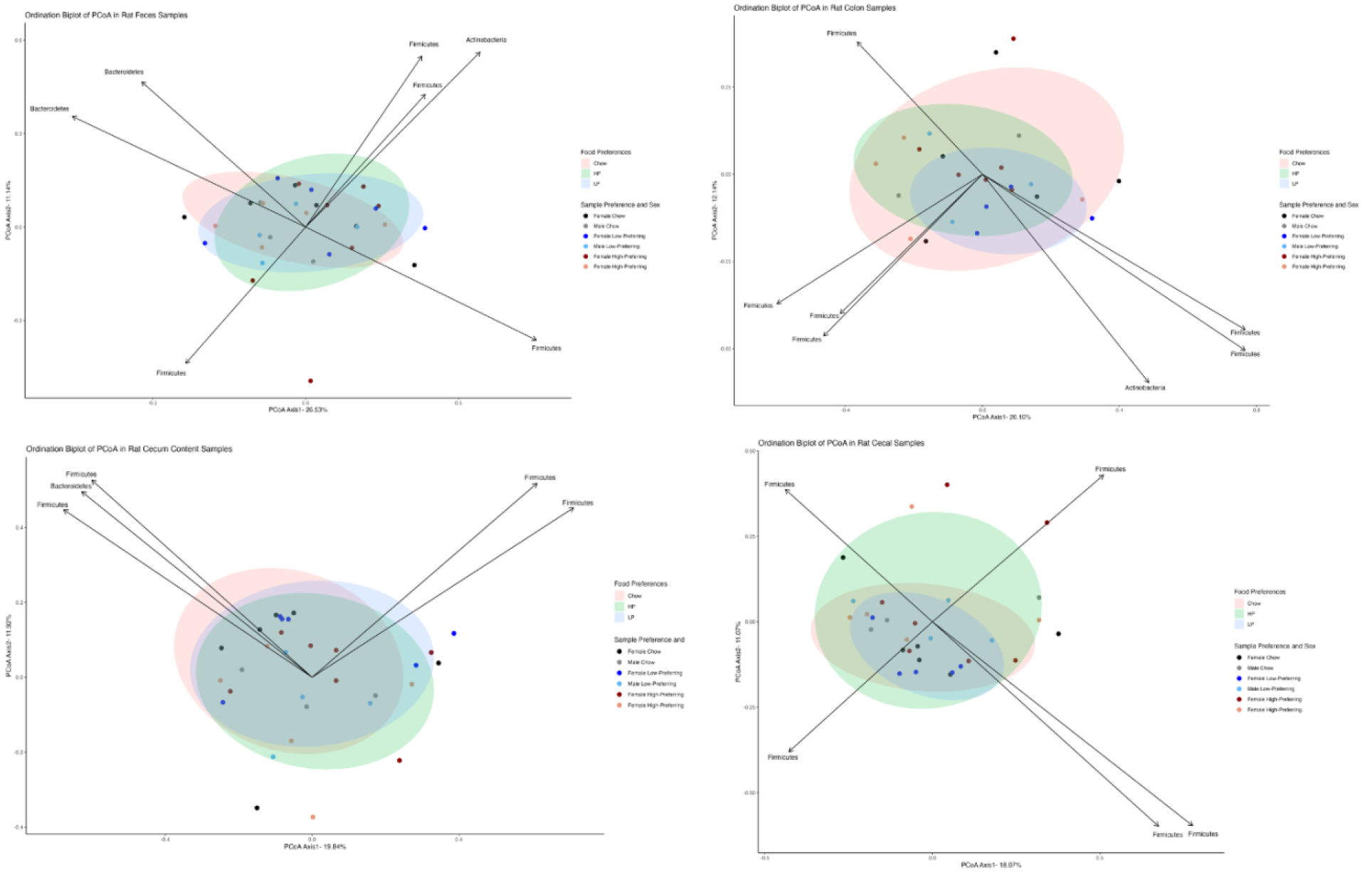
Beta Diversity Measure based on Bray-Curtis distance matrix of the gastrointestinal microbial community showing membership and structure. The percent variation covered by each principal component is indicated in the axis titles. The arrows indicate the top phyla contributors to the community variation. The plot shows an overlap between the food preferences (background colors) and sample preferences based on sex as indicated by the plotted points (OTU-based approach).

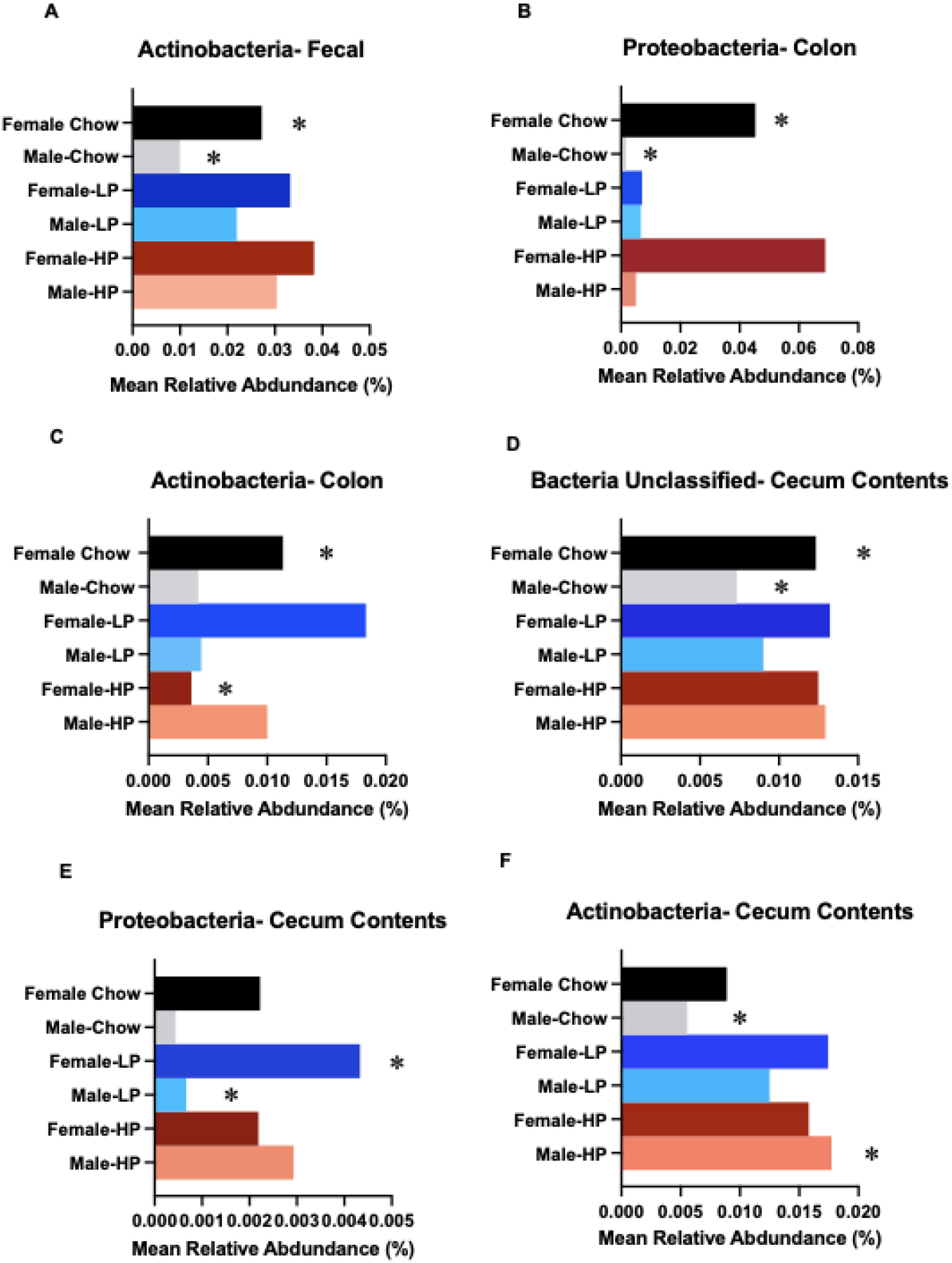

Appendix: The bacterial phyla distribution represents taxa with >2% relative abundance from cecum contents, feces and colon among each feeding behavior group. Significant differences of p<0.05 between groups are shown with (*) next to the bar.

## Discussion

There has been a growing interest in understanding the complex relationship between hosts and their gut microbiota, particularly how this dynamic can impact behavior and overall host health (43,44). A key area of focus that has garnered attention recently is the bidirectional communication system that exists between the brain, gut, and the microbiome (45), which is influenced by factors such as diet, food preference, and sex. Studies have shown that modifying macronutrient and micronutrient intake alters gut microbial composition, which in turn plays a role in shaping feeding behaviors (13, 46). However, the underlying mechanisms driving these interactions, particularly sex specific effects, remain unclear.

To explore these connections, we investigated whether a host’s preference for a high fat, high carbohydrate diet and sex, influence gut microbiota composition. Using an intermittent PF feeding model interspersed with periods of standard chow consumption, we assessed behavioral and microbial changes. Rats were classified into HP and LP groups based on their PF consumption and microbiota differences across sexes were analyzed within these preference groups. Together, these results provide additional insight into the relationship between inherent feeding behavior and gut microbiota composition across distinct regions of the gastrointestinal tract (47).

Analysis of feeding behavior revealed clear preference-dependent differences in food intake across experimental groups. Within the feeding groups, results were consistent with prior studies using a 4-hour feeding paradigm (27). A stronger preference for palatable food and consumed less standard chow consumption was exhibited by HP rats, whereas LP rats consumed more chow during the feeding tests (27, 49). Both HP and LP rats exposed to the palatable diet consumed greater total food than the chow-only group, consistent with previous findings demonstrating increased intake following exposure to a high sucrose/high fat diet (48).

To assess whether intake differences were specific to palatable food exposure, regular chow consumption was evaluated on non-feeding test days, providing a measure of baseline feeding behavior. While one study reported greater chow intake in LP rats compared with HP rats on non-feeding days (27), other studies are consistent with our findings, showing no differences in chow intake between preference groups under these conditions (28). There were also no significant differences in body weight across the groups on non-feeding test days. These findings suggest that preference-based PF is not merely a byproduct of increased caloric intake or body weight in the HP rat groups. Even when intake was standardized relative to body weight, HP rats consistently consumed more palatable food than LP rats, an effect observed in both males and females. Notably, HP rats did not exhibit generalized hyperphagia but instead displayed a distinct, preference-driven feeding pattern when palatable food was available, potentially reflecting gene × environment interactions. These findings suggest that environmental factors can shape feeding behavior even among individuals with similar genetic backgrounds (25). Accordingly, the observed differences in feeding behavior may be associated with shifts in gut microbial communities, raising the possibility that preference-driven intake reflects interactions between genetic predisposition and the microbial environment.

We also examined the relationship between PF consumption and sex differences and found no significant sex-based variations in PF intake in adult rats. Here, we presented findings that suggest no significant sex-dependent difference in PF consumption. Although female HP rats consumed more PF relative to body weight than HP males, this difference was not significant at any time point; a similar pattern was observed in LP rats. These findings are consistent with prior work reporting no sex-specific differences in PF consumption (50). In contrast, other studies have reported sex-dependent differences in food preference (51,52). These discrepancies may reflect differences in experimental paradigms,, including passive (homeostatic consumption) versus active (CPP, behavioral-economics) models. Future studies are needed to clarify how methodological differences contribute to reported sex effects in food preference.

To further examine potential sex × diet × region effects on microbial composition, Metastats analysis was conducted within diet preference groups across multiple gastrointestinal regions. Sex-specific differences were identified within the chow group, with regional variation in relative abundance. In cecal contents, females exhibited lower relative abundance of a shared unidentified taxon compared with males. In the colon, females showed higher relative abundance of Proteobacteria, whereas in fecal samples females exhibited greater relative abundance of Actinobacteria than males. These findings indicate that sex-related differences in microbial composition are region-specific and may vary depending on dietary context. A similar pattern was observed in the cecal contents of LP rats, where females exhibited greater relative abundance of Proteobacteria compared with males. Overall, sex-specific differences in microbial composition were primarily confined to less abundant taxa, whereas higher-level community structure appeared to be driven by dominant phyla, including Firmicutes and Bacteroidetes, and was not significantly influenced by sex.

Sex-based differences in gut microbiota have been hypothesized to arise, at least in part, from hormonal influences (53–56). Empirical evidence supports this framework, demonstrating microbiota shifts following puberty and in response to hormonal manipulations. For example, Yuan et al. (55) reported that although alpha diversity did not differ between males and females in humans, sex-related differences in microbial membership were evident and became more pronounced after puberty. Host sex has also been implicated as a contributing factor in the gut–microbiome–brain axis (57). In rodent models, pre- and postnatal penicillin exposure altered microbiota structure in both sexes; however, only males exhibited reduced anxiety-like behavior and altered sociability, indicating sex-specific behavioral consequences (58). Notably, not all studies report consistent sex effects, underscoring the need for further investigation into developmental and hormonal contributions to microbiome variability (56).

Sex-related differences in microbiota composition may not occur in isolation but instead interact with environmental factors such as diet. There are also multiple lines of evidence supporting the importance of the intersection of diet and sex in shaping microbiome structure. For example, when male and female adult rat populations were given a high fat diet with fish oil, male rats showed an increase in the *Firmicutes/Bacteroidetes* ratio while females showed no comparable shift in gut microbial composition (59). Similarly, diet-driven microbiome shifts in other species---including three spine stickleback fish, mice, and humans have demonstrated sex-dependent patterns (22, 60, 61-64). However, not all studied report significant sex-based differences in response to dietary manipulation, suggesting that the influence of sex may be context-dependent or variably detected across experimental paradigms This finding was noted in the present study for certain taxa, such as *Actinobacteria* whose relative abundance differed according to feeding preference rather than sex. These inconsistencies across studies highlight the need for further research to reconcile methodological inconsistencies and account for developmental stage when evaluating sex effects on microbiome composition (48, 56, 65). While the extent of sex influence on microbiome composition/structure remains debatable, our findings underscore the importance of considering both dietary preference and sex in microbiome research. Supporting this integrative perspective, prior work examining the effects of host genetics, diet, and sex on gut microbiota composition demonstrated that microbial variation can arise from individual factors (e.g., sex or diet), as well as their interactions (e.g., sex × diet) (66).

### Final Thoughts

Dietary modification remains one of the most accessible strategies for altering gut microbiota composition (7), which plays a critical role in host physiology and metabolic health (8-10). Although the extent to which sex independently shapes the gut microbiome remains unresolved, dietary variation clearly influences microbial structure across the gastrointestinal tract. Interpretation of prior published findings has been complicated by methodological heterogeneity and differences in developmental stage across studies. Future studies should adopt integrative designs that examine combined macronutrient exposures, as implemented here, rather than isolated carbohydrate or fat intake. This approach is particularly relevant given that most diets associated with dysbiosis typically involve concurrent elevations in multiple macronutrients rather than one in isolation.

Our findings underscore the importance of characterizing microbiota across multiple GI regions rather than relying on a single sample type. Region-specific assessments provide a more comprehensive understanding of how feeding behavior relates to gut microbial ecology. These findings support a model in which dietary composition, host characteristics, and microbial context interact to shape feeding-related outcomes. Defining how gut microbial communities interact with dietary composition and host factors to influence feeding behavior may ultimately enable targeted microbiome based strategies to improve dietary patterns and metabolic health.

## Notes

### Competing Interest Statement

The authors have declared no competing interest.

### Funding

- U.S. National Science Foundation: IOS-1901546
- National Institutes of Health: DA030444

